# Enspara: Modeling molecular ensembles with scalable data structures and parallel computing

**DOI:** 10.1101/431072

**Authors:** J.R. Porter, M.I. Zimmerman, G.R. Bowman

## Abstract

Markov state models (MSMs) are quantitative models of protein dynamics that are useful for uncovering the structural fluctuations that proteins undergo, as well as the mechanisms of these conformational changes. Given the enormity of conformational space, there has been ongoing interest in identifying a small number of states that capture the essential features of a protein. Generally, this is achieved by making assumptions about the properties of relevant features—for example, that the most important features are those that change slowly. An alternative strategy is to keep as many degrees of freedom as possible and subsequently learn from the model which of the features are most important. In these larger models, however, traditional approaches quickly become computationally intractable. In this paper, we present `enspara`, a library for working with MSMs that provides several novel algorithms and specialized data structures that dramatically improve the scalability of traditional MSM methods. This includes ragged arrays for minimizing memory requirements, MPI-parallelized implementations of compute-intensive operations, and a flexible framework for model estimation.

## I. Introduction

Markov state models (MSMs)^1–3^ are a powerful tool for representing the complexity of dynamics in protein conformational space. They have proven useful both as quantitative models of protein behavior^4–7^ and for producing insights about the mechanism of protein conformational transitions.^8–11^ And, with the rise of special-purpose supercomputers,^12,13^ distributed-computing platforms,^14^ and the dramatic increases in the power of consumer-grade processors (especially GPUs), the size of molecular dynamics (MD) data sets that MSMs are built on have grown in size commensurately.

With the increasing size of MD datasets, there is ongoing and substantial interest in making more tractable models by distilling protein landscapes into a small number of essential states. Typically this is achieved by making assumptions about the relevant features. In particular, existing MSM libraries PyEMMA2^15^ and MSMBuilder3^16–18^ offer state-of-theart, modular components for the newest theoretical developments from the MSM community. These libraries emphasize early conversion to coarse-grained models, particularly through the use of time-lagged independent components analysis (tICA),^19–21^ but also through deep learning^22^ or explicit state-merging.^23,24^ All these approaches merge states that are kinetically close to one another to build a more interpretable model.

Kinetic coarse-graining is effective when the most interesting process is also the slowest, for example when studying folding. However, physiologically-relevant conformational changes also can occur quickly. For example, the opening of druggable cryptic allosteric sites can occur many orders of magnitude faster than the global unfolding process.^25,26^ Thus, for biological questions where the underlying physical chemistry is irreducibly high-dimensional or the features in which it is low-dimensional are not known, building models with a large number of states is an effective strategy for ensuring that important states are not overlooked.

An alternative approach to handling the size of modern MD datasets is to retain the size and high dimensionality, and to learn which features are relevant to the biological question. For example, comparing different sequences’ properties in simulation and in experiments can be one way to learn which features are important. This approach has been successfully leveraged to, for example, understand the determinants of protein stability^7^, enzyme catalysis,^6^ and biochemical properties.^25^ The downside of this approach is that is substantially more computationally demanding, due to the much larger size of both the input features and the resulting model.

In this paper, we present enspara, which implements methods that improve the scalability of the MSM methods. We implement a “ragged array” data structure that enables memory-efficient in-memory handling of data with heterogeneous lengths, and develop tools which use sparse matrices, vastly reduces memory usage of the models themselves while speeding up certain calculations on them. We further introduce clustering methods that can be parallelized across multiple nodes in a supercomputing cluster using MPI, thread-parallelized routines for information-theoretic calculations, and a new framework for rapid experimentation with methods for estimating MSMs.

## II. Results & Discussion

### A. Ragged arrays

The most computation-intensive step in any molecular dynamics-based approach is actually generating the simulation data. One approach to mustering the computation necessary to solve this problem is to harness the power of distributed computing to generate many parallel simulations on many computers. Indeed, one of the points where MSMs excel is in unifying such parallel simulations into a single model. An example of this is the distributed computing project Folding@home.^14^ However, in these scenarios, individual trajectories often substantially differ in their lengths. In Folding@home, the trajectory length distribution shows strong positive skew, with a few trajectories one or more orders of magnitude longer than the median trajectory. Historically, atomic coordinates, as well as features computed on trajectories, have been represented as ‘square’ arrays of *n*_trajectories_ × *n*_timepoints_ × *n*_features_ (or *n*_atoms_ × 3), which assumes uniform trajectory length.^15,27^

To represent non-uniform trajectory lengths, modern implementations of MSM technologies, including the latest version of `MSMBuilder`, use two-dimensional square array with the ‘overhanging’ timepoints filled with a null value. This is also the solution provided by `numpy`,^28^ with its masked array object. While this approach maintains the in-memory arrangement that makes array slicing and indexing fast, it can dramatically inflate the memory footprint of datasets with highly non-uniform length distributions commonly seen in modern molecular dynamics datasets.

In `enspara`, we introduce an implementation of the ragged array, a data structure that relaxes the constraint that the rows in a two-dimensional array be the same length (Fig. 1a). The ragged array maintains an end-to-end concatenated array of rows in memory. When the user requests access to particular elements using a slice or array indices, the object translates these array slices or element coordinates appropriately to the concatenated array, uses these translated coordinates to index into the concatenated array, and then reshapes the data appropriately and returns it to the user. On trajectory length distributions as described, the ragged array scales much better than the padded square array (Fig. 1b).

**FIG. 1.**
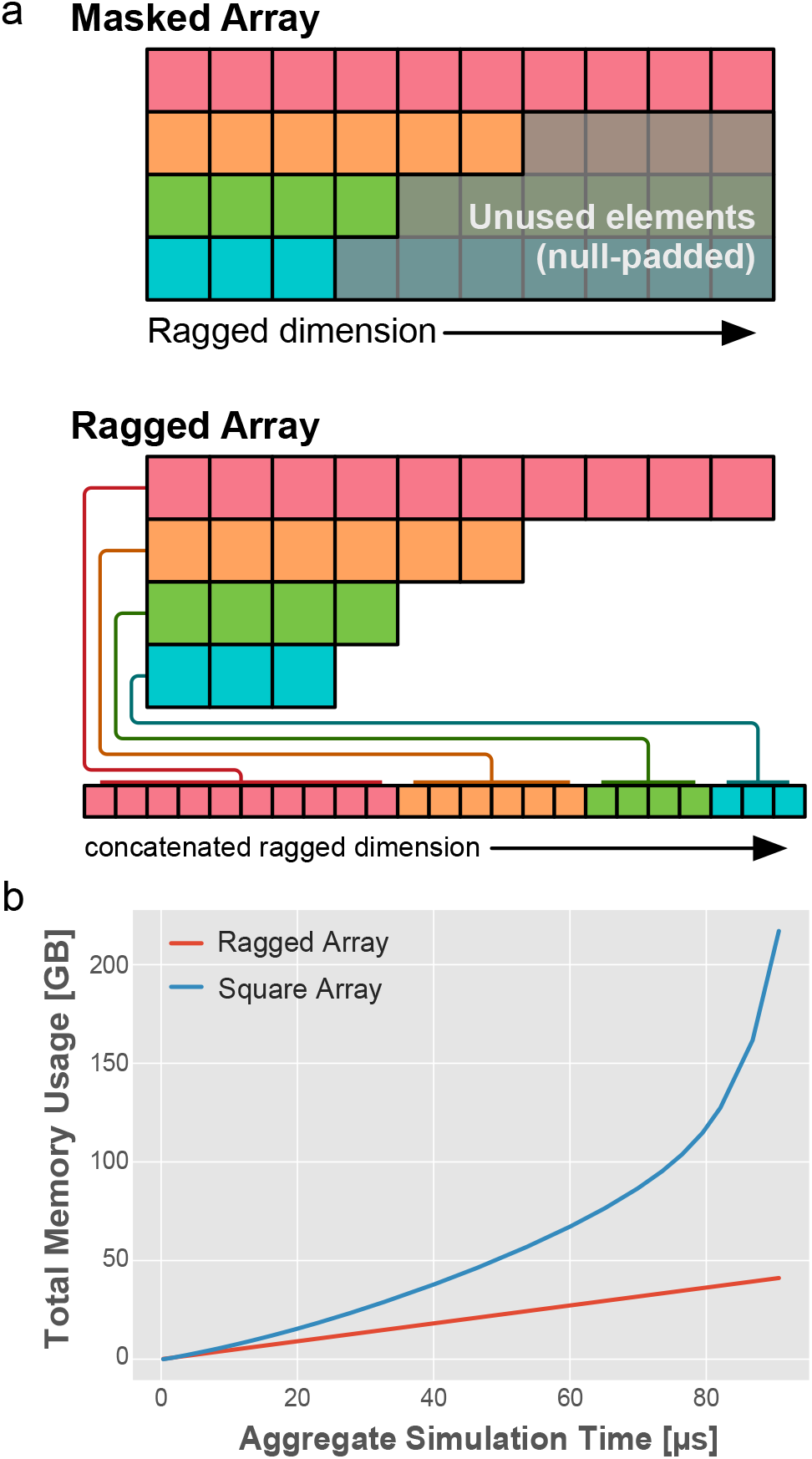
Ragged arrays compactly store non-uniform length data in memory. **a**, A schematic comparison between the memory footprint of a masked, uniform array and our implementation of the ragged array interface. In the masked array, rows of length lower than the longest row are padded with additional, null-valued elements to preserve the uniformity of the array. In the ragged array, however, rows are stored concatenated and memory is not expended. **b**, A plot of memory used by traditional and ragged arrays as a function of aggregate simulation time as trajectories of increasing length are added from a previously-published Folding@home dataset.

Our implementation of the ragged array is a subclass of `numpy’s ndarray` class. Consequently, most functions that work on ndarrays also function correctly on our ragged array class. This allows users to leverage the sophisticated slicing and indexing operations available in the traditional two-dimensional array without sacrificing large portions of memory to null-filled elements.

### B. SIMD clustering using MPI

One of the most expensive and worst-scaling steps in the Markov state model construction processes is clustering. The most popular clustering algorithms for use in the MSM community are *k*-means^29^ (generally composed of *k*-means++^30^ initialization and Lloyd’s algorithm^31^ for refinement) for featurized data, and *k*-hybrid^17^ (composed of *k*-centers^32^ initialization and *k*-medoids^33^ refinement) for raw atomic coordinates. Both of these algorithms scale roughly with *O*(*nkdi*), where *n* is the number of observations, *d* is the number of features per observation, *k* is the number of desired cluster centers, and *i* is the number of iterations required to converge. Unfortunately, with the possible exception of *i*, these numbers are all generally very large. As discussed above (Section II C), the number of clusters *k* must be large for some problems, proteins are intrinsically high-dimensional objects (*i.e*. high *d*), and the increasing speed of simulation calculations^34^ has increased the number of timepoints that must be clustered, *n*, into the millions.

To address the poor scaling of clustering, the MSM community has developed a number of approaches to managing this problem. One approach is to reduce the number of observations by subsampling data^27^ so that only every *n* th frame is used. Another approach is to reduce the number of features by including only certain atoms (as in Zimmerman *et al*.^7^, Schwantes and Pande^35^, Bowman and Geissler^36^), using a dimensionality reduction algorithm like principal components analysis (PCA),^37,38^ or creating a hand-tuned set of order parameters (*e.g*. specific, relevant pairwise atomic distances). Yet a third approach is to use tICA^19,20^ as a dimensionality reduction, which has the benefit of reducing both the number of features and the number of clusters needed to satisfy the Markov assumption, but has the disadvantage that it may obscure important fast motions and can be sensitive to hyperparameters (in particular the lag time).^19^

As an alternative or complimentary approach to preprocessing data to reduce input size is to parallelize the clustering algorithms themselves so that many hundreds, rather than many tens, of cores can be simultaneously utilized. Message Passing Interface (MPI)^39^ is a parallel computing framework that enables communication between computers that are connected by low-latency, high-reliability computer networks, like those commonly encountered in academic cluster computing environments. This approach to interprocess communication has enabled numerous successful parallel applications including molecular dynamics codes like GROMACS^40,41^ (among many others). This approach to interprocess communication allows information to be shared easily across a network between an arbitrary number of distinct computers. Thus, for a successfully MPI-parallelized program, the amount of main memory and number of cores available is increased from what can be fit into one computer to what that can be fit into one supercomputing cluster—a difference of one or two dozens of processors to hundreds of processors. However, because interprocess communication is potentially many orders of magnitude slower than, for example, in thread-parallelization, single-core algorithms must generally be adjusted to scale well under these constraints.

In this work, we present low-communication, same-instruction-multiple-data (SIMD) variants of a clustering algorithms that are popular in the the MSM community, *k*-centers, *k*-medoids, and *k*-hybrid.^17^ Specifically, data—atomic coordinates and distances between coordinates and medoids—are distributed between parallel processes which can reside on separate computers, allowing more data to be held in main memory, and allowing more processors *in toto* to be brought to bear on the data.

The *k*-centers initialization algorithm^32^ repeatedly computes the distance of all points to a particular point, and then identifies the maximum distance amongst all distances computed this way. This introduces the need for communication to (1) distribute the point to which distances will computed and (2) collectively identify which distance is largest. (1) is solved trivially by the MPI `scatter` directive and (2) is solved using computing local maxima and then distributing these maxima with MPI `allgather`. Implementation details of *k*-medoids are somewhat more complex but follow a similar pattern. The performance characteristics of this implementation as a function of data input size is plotted in Fig. 2a and b, which show marked decreases in runtime as additional computers are added to the computation. The *k*-hybrid algorithm simply performs *k*-centers followed by *k*-medoids.

**FIG. 2.**
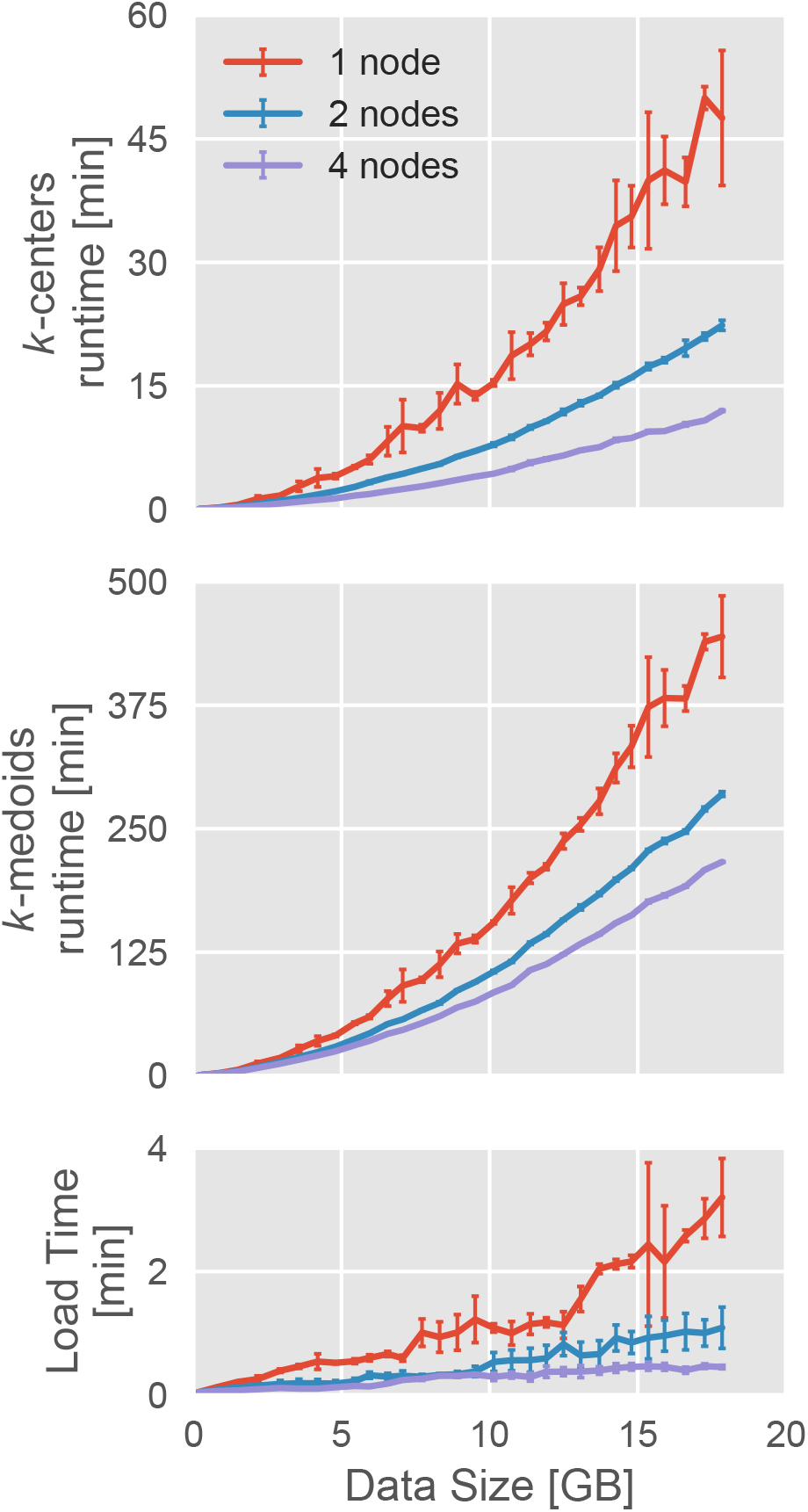
SIMD reformulation of clustering algorithms allows greater scaling. **a**, The runtime of the parallelized *k*-centers code as a function of data input size. **b**, The runtime of the parallelized *k*-medoids code as a function of data input size. **c**, The load time of the parallel code as a function of input data size. Points represent the average and error bars the standard deviation across three trials.

A further advantage of a parallelized algorithm is that, if configured correctly, it can also decrease load times. In the traditional HPC environment used in many academic settings, data typically resides on a single central, “head” node and it is distributed to “worker” nodes via a network file system (NFS). The NFS can transfer data to any particular worker node only as quickly as the network allows, which is generally orders of magnitude slower than the rate at which it can be loaded from disk into memory. However, if network topology allows nodes to independently communicate with the head node, the network bottleneck is reduced or removed and load times can be substantially decreased, as shown in Fig. 2c. While load times do not dominate the overall runtime of the algorithms we discuss here, loading data represents a very frequent failure point in misconfiguration, and so short load times are immensely helpful from a usability perspective.

### C. Sparse matrix integration

Building a Markov state model with tens of thousands of states presents some methodological challenges. One of these is the representation of the transition counts and transition probability matrices. Most straightforwardly, this is achieved using dense arrays, such as the `array` or `matrix` classes available in numpy, and this is the strategy employed by extant MSM softwares, `MSMBuilder3`^17,18,27^ and `PyEMMA`^15^. The problem with this representation is that the memory usage of these matrices grows with the square of the number of states in the model. To make matters worse, the computational cost of the eigendecomposition that is typically required to calculate a model’s stationary distribution (equilibrium probabilities) and principal relaxation modes grows with the cube of the number of elements in the matrix.^42^

To address the computational challenges posed by traditional arrays, enspara has been engineered to support sparse arrays wherever possible. Sparse arrays have been supported by `MSMBuilder` in the past, but were dropped with version 3. PyEMMA also makes heavy use of dense arrays, although there is some support for sparse arrays. Sparse arrays, rather than growing strictly with the square of the number of states, grows linearly in the number of nonzero elements in the array. In the worst case, where every element of the transition counts matrix is non-zero (*i.e*. every possible transition between pairs of states is observed) this becomes the dense case. However, this is very unusual: the number of observed transitions is generally several orders of magnitude smaller than the number of possible transitions (Fig. 3a). By implementing routines that support `scipy`’s sparse matrices, it becomes possible to keep much larger Markov state models in memory (Fig. 3b) and analyze those models much more quickly (Fig. 3c).

**FIG. 3.**
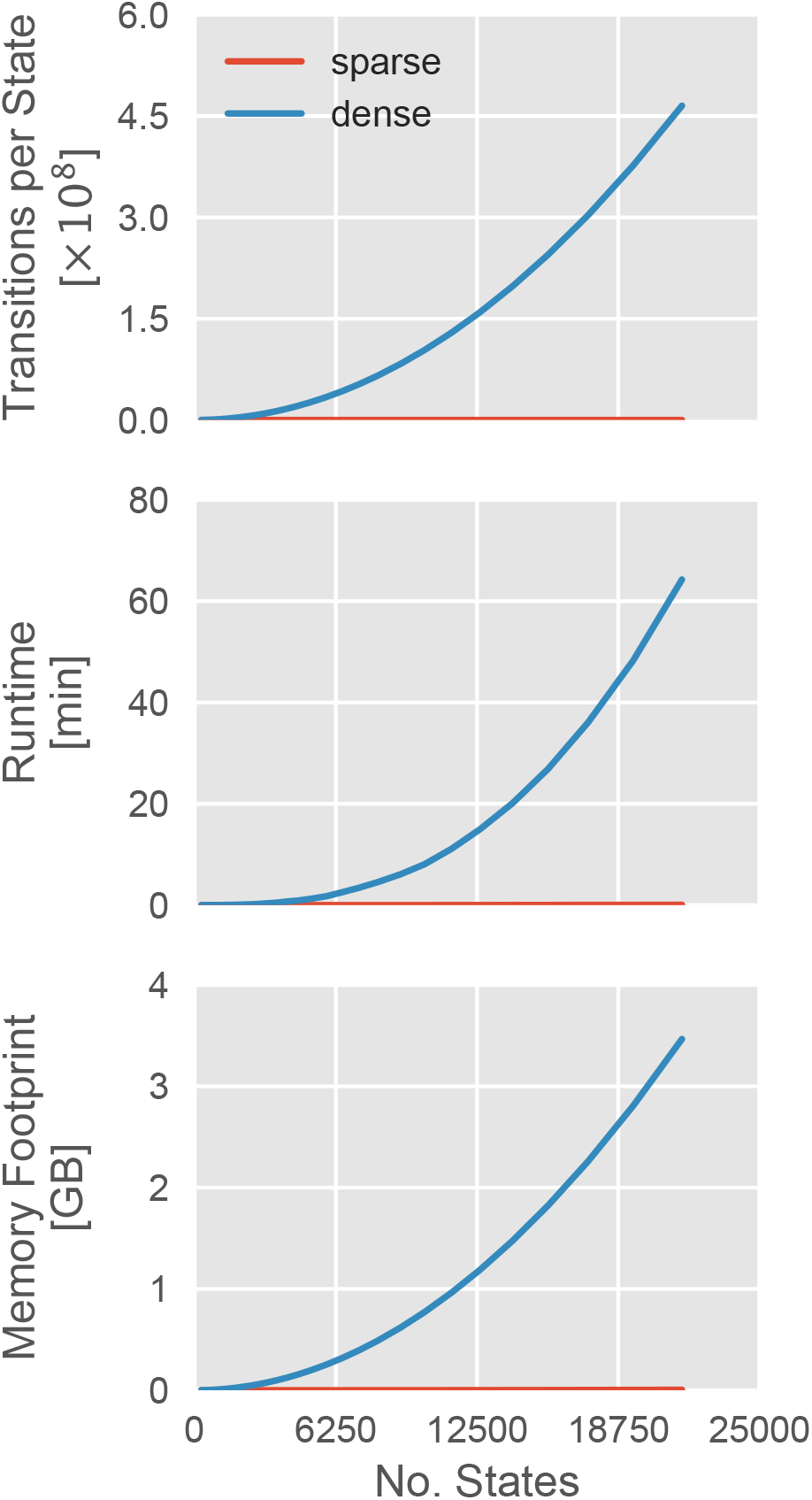
The performance characteristics of sparse and dense matrices representing the same MSM. **a**, The runtime of an eigendecomposition as a function of the number of states in a model. **b**, The memory footprint of the transition probability matrix as a function of the number of states in a model. **c**, The number of transitions per state in a transition counts matrix as a function of the number of states in the model.

### D. Fast and MSM-ready information theory routines

Recent work^43–45^ has demonstrated the usefulness of information theory, and mutual information (MI) in particular, for identifying and understanding the salient features of conformational ensembles. MI is a nonlinear measurement of the statistical non-independance of two random variables. MI is given by

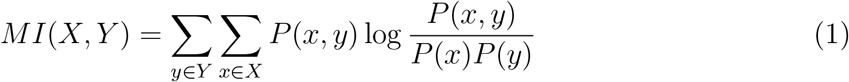

where *P*(*x*) is the probability that random variable *X* takes on value *x, P* (*y*) is the probability that random variable *Y* takes on value *y*, and *P* (*x, y*) is the joint probability that random variable *X* takes on value *x* and that random variable *Y* takes on value *y*.

Historically, the joint distribution *P* (*x, y*) is estimated by counting the number of times that combination of features appeared in each frame.^43,44^ This computation can become a bottleneck when it must be computed over hundreds or thousands of different features and for datasets with hundreds of thousands or millions of observations. This is because it is highly iterative—which is notoriously slow in many higher-level programming languages like python or Matlab—and because the number of joint distributions that must be calculated grows with the square of the number of features to be tracked. Consequently, in the worst case, this involves examining every frame of a trajectory *n*^2^ times, where *n* is the number of random variables of interest.

In `enspara`, we take two overlapping approaches to address the problem of the poor scalability of pairwise MI calculations. The first approach is to use the joint distribution implied by the equilibrium probabilities of a Markov state model, rather than by counting co-occurrences from full trajectories. Specifically, the joint probability *P* (*x, y*) is estimated by 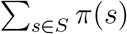, where *S* is the set of states where *x* = *X* and *y* = *Y* and *π*(*s*) is the equilibrium probability of state *s* from the MSM. This works by reducing the number of individual observations, usually by orders of magnitude. Existing codes^44,46^ either do not provide the option to compute MI with weighted observations or require a specific object-based framework to do so.^47^

Our second approach is to implement a fast joint counts calculation routine. This routine is both thread-parallelized and much faster than the equivalent numpy routine even on a single core. This approach is needed because, in some cases (*e.g*. Singh and Bowman^44^), information from a Markov state model cannot be trivially substituted for frame-by-frame calculations. To address this case, we also implement a function using cython^48^ and OpenMP^49^ that takes a trajectory of *n* features and returns a four-dimensional joint counts array with dimension *n* × *n* × *s_n_* × *s_n_*, where *s_n_* is the number of values each feature *n* can take on. The value of returning this four-dimensional joint counts matrix is that it renders the problem embarrassingly parallel in the number of trajectories: this function can be run on each trajectory totally independently, and the resulting joint counts matrices can be summed before being normalized to compute joint probabilities. We recommend combining this with a pipelining software like Jug.^50^

Additionally, in this package, we include a reference implementation of Correlation of All Rotameric and Dynamical States framework (CARDS).^44^ In brief, this method takes a series of molecular dynamics trajectories and computes the mutual information (MI) between all pairs of dihedral angle rotameric states, and between all pairs of dihedral angle order/disorder states. A dihedral angle is considered disordered if it frequently hops between rotameric states. This implementation parallelizes across cores on a single machine using the thread-parallelization described in Section II D.

### E. Fast and flexible model fitting

Estimating a transition probability matrix from observed state transitions is a crucial step in building an MSM, yet there is not a uniform procedure for accomplishing this that works in all cases. Many different estimators exist for generating transition probabilities, and more are in active development. Perhaps the simplest procedure to estimate the transition probability matrix, *T*, is to row-normalize the transition count matrix, *C*,

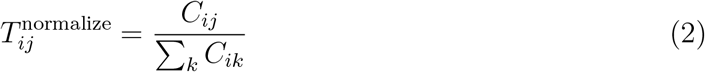

where *T_ij_* is the probability of observing a transition from state *i* to *j* and *C_ij_* is the number of times such a transition was observed. While this method is simple, it can and often does generate a non-ergodic state space. In an effort address this difficulty and to condition the MSM to be well-behaved, one can include an additional pseudocount *ĉ* before estimation,

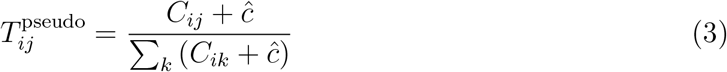

which ensures ergodicity.^51^ A more dramatic conditioning comes forcing the counts matrix to obey detailed balance by averaging forward and reverse transitions:

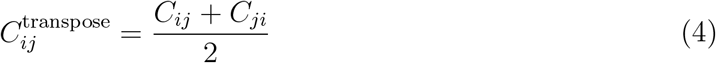

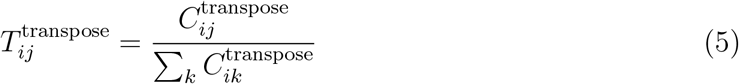

Yet a third proposed way of estimating an MSM is to find the maximum likelihood estimate for *T* subject to the constraint that it satisfies detailed balance.^27,52^ Framed as a Bayesian inference, the transition probabilities are solved as the most likely given a transition counts matrix, such that,

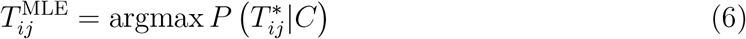

Additionally, there exist more sophisticated schemes of estimation, such as those that draw on inspiration from observable operator models,^53^ and projected MSMs.^54^ While it is beyond the scope of this article to review this area of study in exhaustive detail, we hope these few examples demonstrate the variety and importance of estimators. This poses a major challenge to writing a framework that can readily estimate a transition probability matrix; estimators are an active area of research, and a flexible framework that allows users to quickly modify an existing estimator or try a new one would be of great utility.

To facilitate experimentation with various methods for fitting an MSM, we have developed an API that separates the responsibility of fitting an MSM from the MSM itself. We call the methods that take a transition counts matrix and return transition and equilibrium probabilities `builders`. These built-in functions, along with our MSM object can be used to quickly fit an MSM using commonly-used approaches (Fig. 4a). Alternatively, for users who wish to slightly modify existing MSM estimation methods, the function-level interface provides fine-grained control over the steps in fitting an MSM (Fig. 4b). Finally, for users who wish to prototype entirely new MSM estimation methods, any function or callable object is accepted as a `builder`, as long as it accepts a transition counts matrix *C* as input and returns a 2-tuple of transition probabilities and equilibrium probabilities.

**FIG. 4.**
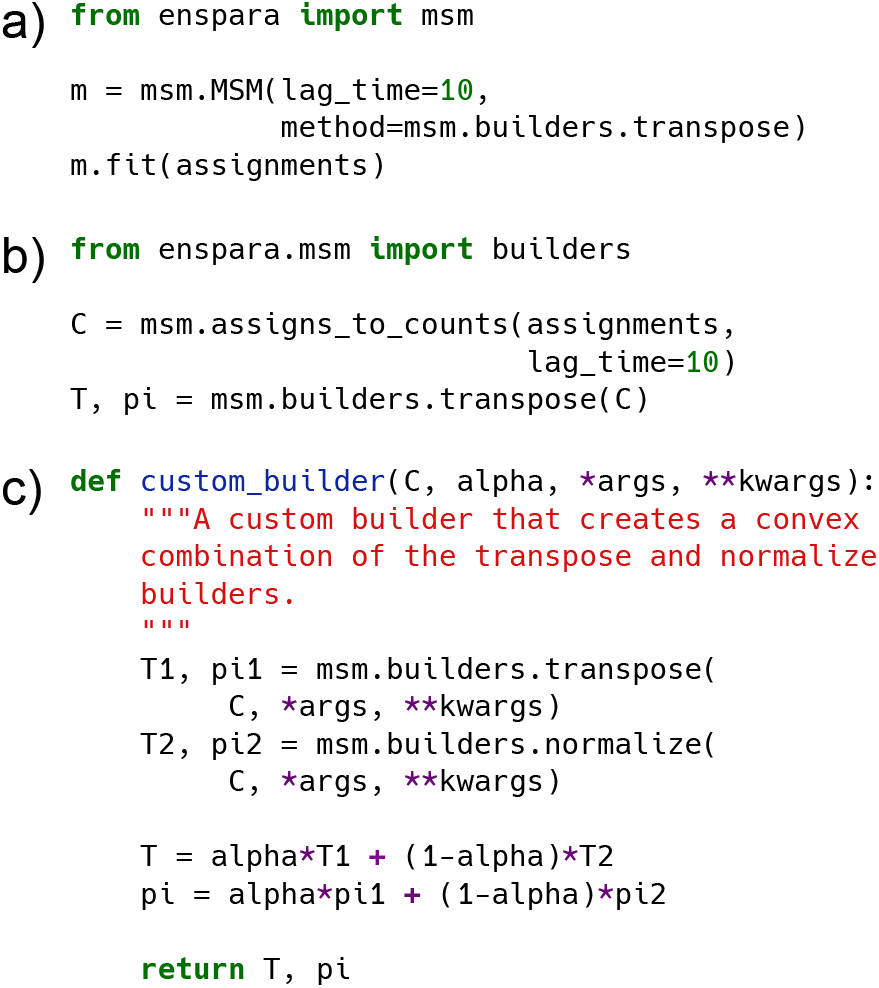
**a**, An example usage of the high-level, object-based API to fit a Markov state model. **b**, An example usage of `enspara`’s low-level, function-based API to fit a Markov state model. **c**, A custom method that fits a Markov state model and is interoperable with `enspara`’s existing API.

## III. Methods

### A. Libraries and Hardware

Eigenvector/eigenvalue decomposition experiments were performed on a Ubuntu 16.04.5 (xenial) workstation with an Intel i7–5820K CPU @ 3.30GHz (12 cores) with 32GB of RAM using SciPy version 1.1.0 and numpy 1.13.3. Probabilities were represented as 8-byte floating point numbers.

Thread parallelization experiments were performed on the same hardware using OpenMP 4.0 (2013.07) with gcc 5.4.0 (2016.06.09) and cython 0.26 in Python 3.6.0, distributed by Continuum Analytics in conda 4.5.11.

Clustering scaling experiments were performed on identical computers running CentOS Linux release 7.3.1611 (Core) with Intel Xeon E5–2697 v2 CPUs @ 2.70GHz and 64 GB of RAM linked to a head node with two Intel 10-Gigabit X540-AT2 ethernet adaptors and nfs-utils 1.3.0. We used the `mpi4py`^55–57^ and Python 3.6.0 with Open MPI 2.0.2. Clustering used as a distance metric the RMSD function provided in the MDTraj 1.9.1.^58^

### B. Simulation data

For example simulation data, we used a previously-published 90.5 *μ*s TEM-1 *β*–lactamase dataset. As described previously,^10^ simulations were run at 300 K with the GROMACS software package^40,41^ using the Amber03 force field^59^ and TIP3P^60^ explicit solvent. Data was generated using the Folding@home distributed computing platform.^14^

## IV. Conclusion

In this work, we have presented `enspara`, a library for building Markov state models at scale. We introduced a `numpy`-compatible implementation of the ragged array, which dramatically improved the memory footprint of MSM-associated data. We developed a low-communication, parallelized version of the classic *k*-centers and *k*-medoids clustering algorithms, which simultaneously reduce runtime and load time while vastly increasing the ceiling on memory use for those algorithms by allowing execution on multiple computers simultaneously. `enspara` also has turn-key sparse matrix usage. Finally, we implement a function-based API for MSM estimators that greatly increases the flexibility of MSM estimation to enable rapid experimentation with different methods of fitting. Taken together, these features make enspara the ideal choice of MSM library for many-state, large-data MSM construction and analysis.

## Acknowledgments

We are grateful to the Folding@home users for computing resources. This work was funded by National Institutes of Health Grants R01GM12400701, U19AI109664, and T32GM02700, as well as by the National Science Foundation CAREER Award MCB-1552471. G.R.B. holds a Career Award at the Scientific Interface from the Burroughs Wellcome Fund and a Packard Fellowship for Science and Engineering from The David & Lucile Packard Foundation. M.I.Z. holds a Monsanto Graduate Fellowship and a Center for Biological Systems Engineering Fellowship.

